# Tank Fouling Community Enhances Coral Microfragment Growth

**DOI:** 10.1101/2022.07.04.498770

**Authors:** Chris Page, Riley Perry, Claire Lager, Jonathan Daly, Jessica Bouwmeester, E. Michael Henley, Mary Hagedorn

## Abstract

Anthropogenic stressors threaten reefs worldwide and natural *in situ* coral reproduction may be inadequate to meet this challenge. Land-based culture can provide increased coral growth, especially with microfragments. We tested whether culture methods using different algal fouling communities could improve the growth and health metrics of microfragments of the Hawaiian coral, *Porites compressa*. Culture method fouling communities were: 1) similar to a reef environment (Mini Reef); 2) clean tanks managed to promote crustose coralline algae (Clean Start); and, 3) tanks curated beforehand with poorly-competing algae (Green Film) assessed in winter and summer months. The Green Film method during the winter produced the fastest microfragment mean growth at 28 days until the first row of new polyps developed and the highest tank metric scores. Time efficient, standardized methods for land-based culture designed to maximize growth and production of coral fragments will contribute considerably to the success of large scale restoration efforts.

## Introduction

Coral reefs worldwide are dying due to widespread local and global anthropogenic stressors (Hughes et al. 2003; Richmond et al. 2018). This is occurring so rapidly that many coral species may face severe extinction risk by 2100 (Hoegh-Guldberg et al. 2017; Logan et al. 2021). In light of this reef crisis, scalable mechanisms to hasten the recovery of impacted reefs and the adaptation of corals to global stressors have been proposed (National Academies of Sciences & Medicine 2019). However, focusing on *in situ* coral propagation strategies alone may be inadequate to meet this challenge, as producing survivable early life stages of many key species remains an inefficient and slow process (dela Cruz et al. 2015; Krumholz et al. 2010).

Recent studies with land-based culture techniques have led to increased growth and survival of young coral (Forsman et al. 2015; Hagedorn et al. 2021; Page et al. 2018), and have been proposed to augment *in situ* processes via large-scale restoration projects such as Mission Iconic Reef in Florida (USA) and the Reef Restoration and Adaptation Program in Australia. Despite these successes, outcomes between facilities are often variable due in large part to differing coral culture conditions (Hagedorn et al. 2021; Petersen et al. 2006). This variable performance will greatly impact the utility and scale of land-based culture efforts if prolific, replicable, and transferrable techniques cannot be identified and implemented.

Algal fouling can be a substantial source of competition and mortality for cultured corals. Algae compete with corals via direct overgrowth, allelopathy, and by facilitating microbe proliferation (Kuffner and Paul 2004, Vermeij et al. 2009). Generally, to combat these effects aquarists aim to grow coral by creating a mature reef mesocosm or mini reef, e.g., (Craggs et al. 2019). Though successful, each mini reef system can produce a unique mixture of noxious and benign phototrophic fouling organisms over time, making them difficult to study and replicate. We hypothesized that quickly establishing a simple, poorly-competing, fouling community during coral culture would lead to increased microfragment growth. To test this hypothesis, we used uniformly-sized microfragments from the Hawaiian coral, *Porites compressa*, and compared their growth and health with identified fouling metrics over two years using three sequential culture methods: 1) a mini reef mesocosm (Mini Reef); 2) clean tanks managed to promote Crustose Coralline Algae (CCA) and, 3) tanks curated beforehand with poorly-competing green algae (Green Film). The results of this study can be used to improve the replicability of land-based coral culture techniques.

## Methods

### Collection of coral

*Porites compressa* colonies (15 cm) were collected from reefs in Kaneohe Bay, Oahu, HI (USA) in accordance with Special Activity Permit numbers 2020-25, 2021-33, and 2022-22 from the Hawaii Department of Land and Natural Resources. Care was taken to collect colonies at different locations on various reefs to avoid collecting clones of the same genotype.

### General care of coral, microfragmentation, and husbandry

The general care of coral, process of microfragmentation and establishing each of the three culture methods can be found in Supplemental Methods. Briefly, coral colonies were cut into 30 uniformly-sized microfragments (1.0 × 0.75 cm) and glued onto a plastic sheet supported by a plexiglass plate (Fig 1). Microfragments were then grown in our husbandry system using three sequential culture methods where the fouling communities were: 1) a reef mesocosm (Mini Reef; February to May 2020), a culture technique used by many laboratories and aquaria; 2) clean tanks managed to try and promote CCA settlement as described in Hagedorn et al (2021) (Clean Start; April 2020 to September 2020); and, 3) tanks curated beforehand with a poorly-competing green algae (Green Film; October 2020 to June 2021), a new culture method devised for this work (Fig 1S).

**Fig. 1.**
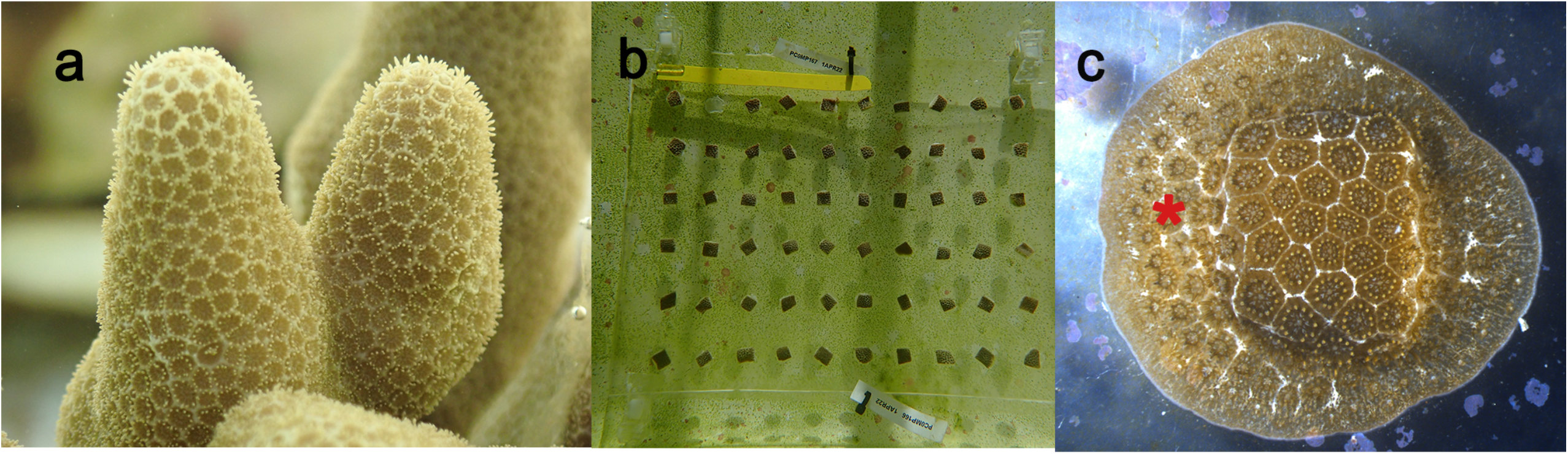
Images of *Porites compressa* used in these experiments **a**. Colonies of *P. compressa* before cutting. **b**. The microfragment array after cutting and gluing the 0.75 × 1 cm pieces of *P. compressa* onto a plastic sheet clipped to a plexiglass plate for support. Note, the conspicuous green film algae showing through the plexiglass plate. Inset shows details of the phytoplankton cells (∼10µm) that make up the green film. **c**. A growing microfragment that has three to four rows of new polyps (*) sheeting out around the cut microfragment. We defined the onset of growth as the date when one ring of new polyps surrounds the microfragment.

These methods were developed and used sequentially as part of another microfragment experiment that required rapid and replicable tissue growth. A new method was therefore adopted once it became apparent that the current method was not suitable, and consequently the Mini Reef and Clean Start methods were only utilized for 3 and 5 months respectively.

### Microfragment Growth Rate and Qualitative Metrics

#### Microfragment Growth Rate

For this study, the onset of new polyps signaled that the coral had healed, resumed calcification, and was in a healthy, robust growth phase. To understand how microfragment growth differed amongst the three culture methods, for each sampled microfragment the days from the microfragment cut date until it produced the first ring of new polyps was recorded and compared (Mini Reef: n=3 genets, 7 microfragments; Clean Start: n=19 genets, 106 microfragments; Green Film Summer: n=25 genets, 196 microfragments; Green Film Winter: n=23 genets, 139 microfragments).

#### Tank and Plate Scores and Microfragment Health Score

In addition to growth, we developed a suite of qualitative metrics to understand the health of the tank, the plate and the microfragments (that had achieved 1 row of new polyps) in relationship to fouling. During the Clean Start and Green Film methods qualitative metrics composed of 3 categories were assessed: 1) CCA health: whether CCA was growing or dying in the tank; 2) CCA recruitment: the ratio of new CCA growing in the tank compared to other competing algae; and, 3) nuisance algae coverage: how much and what type of algae was growing throughout the tank (see Table 1S for scoring criteria). Fouling was visually assessed on all surfaces of the tank including any added substrates (i.e., heaters, air stones, snail cages, etc.) at the meter scale. These values were then totaled to determine a score for each metric. The higher the score, the healthier the tank or plate was considered to be. The Clean Start method was only used during the summer as it became supplanted by the Green Film method. Data collected during the green film method was divided into values collected during winter months (Dec 1 to Mar 30) and all other months to determine if season effected outcome. As the qualitative metrics were developed in the early stages of the Clean Start method they were not available for application to the earlier Mini Reef approach.

Health metrics for individual microfragments were assessed and compared, because each microfragment experienced differing conditions based on their position among and within culture tanks, using three metrics: 1) polyp extension (i.e., the percentage of microfragments on the plate that had their polyps extended); 2) change in coloration/ tissue recession: the number of microfragments that had paled or had tissue recession; and, 3) new tissue integrity: the thickness and color of the tissue surrounding the microfragment and its rate of advancement (see Table 2S). The higher the score, the healthier the microfragment was considered to be. The number of genotypes in each method was Clean Start Summer: n=17 genets, 162 microfragments; Green Film Summer: n=29 genets, 244 microfragments; Green Film Winter: n=21 genets, 119 microfragments.

### Statistical Methods

To analyze the difference in the means for the microfragment growth rate in response to each of the three culture methods, a Kruskal-Wallis non-parametric test was performed on the number of days needed for growth, followed by a Dunn’s Multiple Comparison *post-hoc* test. To analyze the difference in the means for the fouling and health metrics, the scores were analyzed with a non-parametric Kruskal-Wallis test with a Dunn’s Multiple Comparison *Posthoc* Test. Statistical analyses were performed in Graph Pad Prism 9.0 (San Diego, CA) and R (R Core Team 2019). All errors are represented as standard error of the mean, unless otherwise stated.

## Results

Growth was defined as the total number of days from initial cut to generation of the first ring of polyps (Fig. 1 and 2). The Green Film method overall produced the most rapid growth especially during winter months (28.3±0.5 days; Kruskal-Wallis χ^2^_(3)_ = 162.5, p < 2×10^−16^ and Dunn *post-hoc* test; Fig. 2) with a 3.3-fold increase in growth rate compared to the Mini Reef method (91.9±3.1 days). For experimental and restoration purposes, rapid healing and growth from uniformly cut microfragments will be an important element toward producing material as quickly as possible in all coral species.

**Fig. 2.**
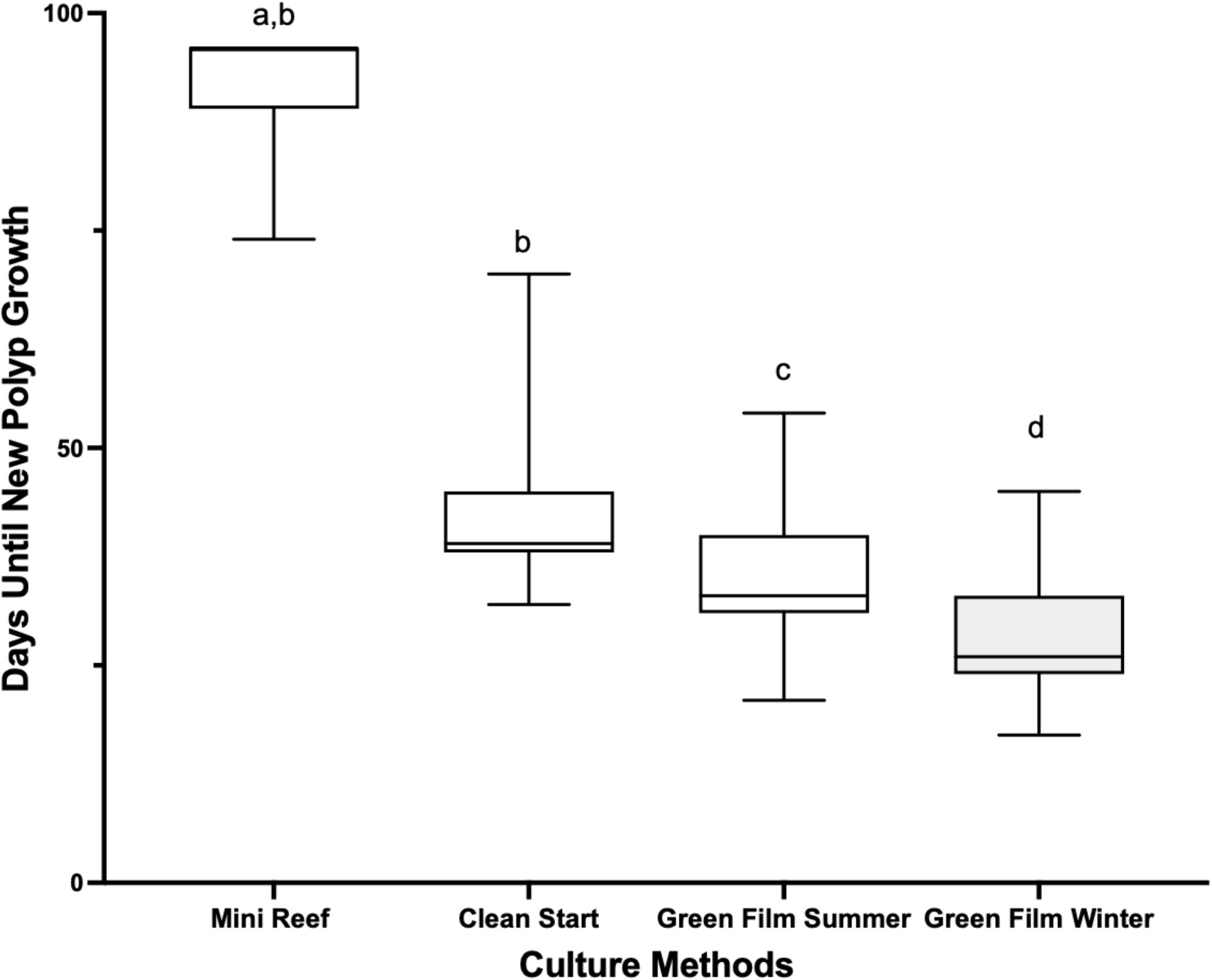
Mean growth rates of the *P. compressa* microfragments under three culture methods with two seasons for Green Film The time until the first row of new polyps was measured for each culture method and season. The Mini Reef (7 microfragments; n=3) has a mean growth time of 91.9±3.1 days, whereas the Clean Start (106 microfragments; n=19 had a mean growth time of 40.9±0.6 days), Green Film Summer (196 microfragments; n=25 had a mean growth time of 97±4.5 days), and Green Film Winter (139 microfragments; n=23 had a mean growth time of 28.3±0.5 days) reducing the time to first growth 2.3, 2.6 and 3.3-fold, respectively. The box represents the central 50% of the data distribution, the bar in the box represents the median, the whiskers represent the remaining 50% data spread. Different letters above plots indicate significant differences (α = 0.05), as determined by Dunn’s Multiple Comparison *posthoc* test.

Similarly, the Green Film method conducted during the winter produced the highest mean tank, plate fouling, and microfragment health scores (Kruskal-Wallis (tank) χ^2^_(2)_ = 106.9, p < 2×10^−16^, (plate) χ^2^_(2)_ = 80.3, p < 2×10^−16^, (microfragment) χ^2^_(2)_ = 27.7, p = 1×10^−6^, and Dunn *posthoc* tests; Fig. 3; Table 1S and 2S for metrics definitions).

**Fig. 3.**
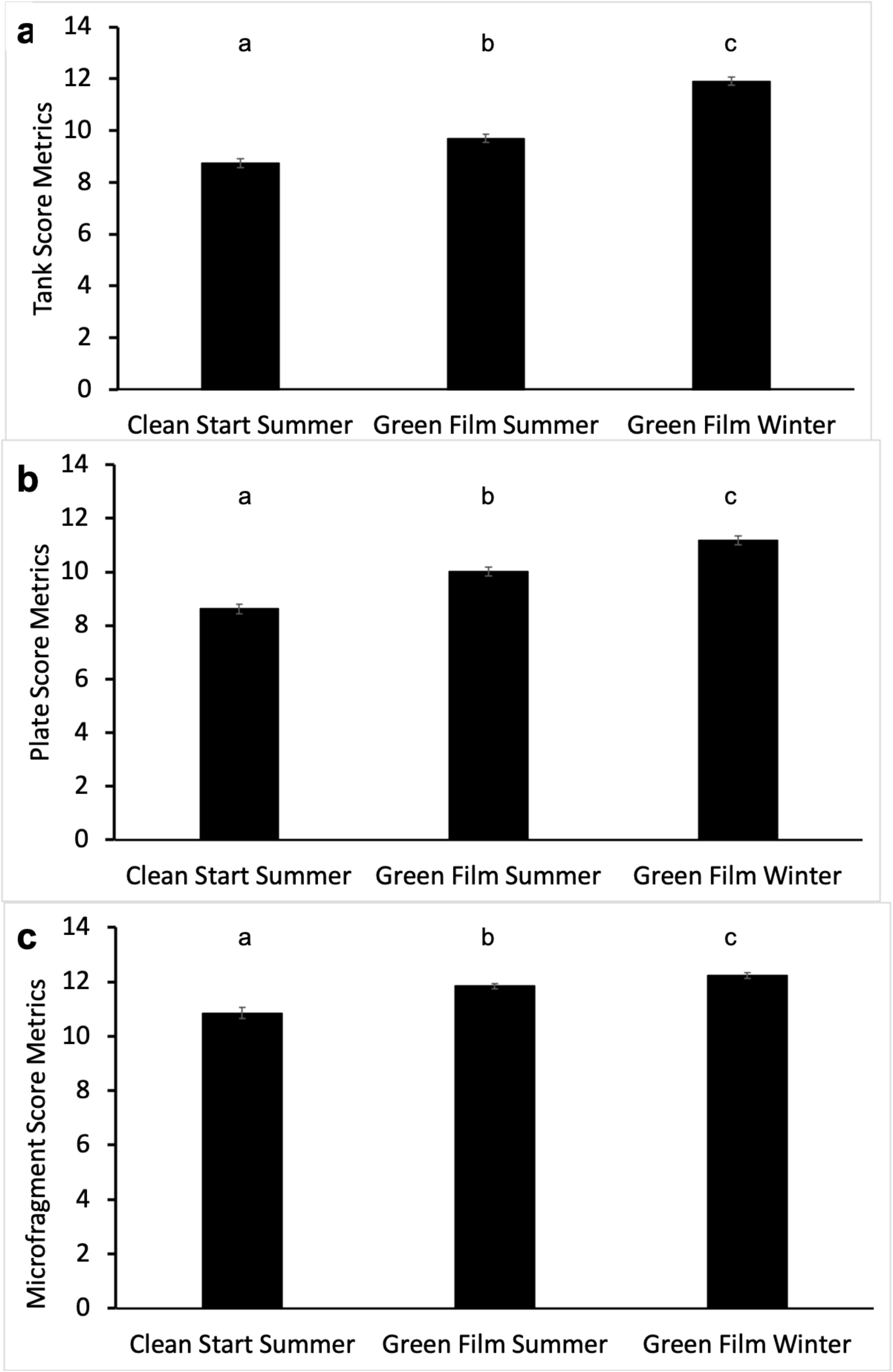
Fouling and health metrics versus culture methods The tanks (a) and plates (b) in various tanks in different culture methods were scored by assessing tank fouling metrics; 1) CCA health; 2) CCA recruitment; and, 3) nuisance algae coverage. The microfragments in these tanks were scored by assessing: 1) polyp extension; 2) change in coloration/ tissue recession; and, 3) new tissue integrity. The Green Film Winter had the highest overall mean metric scores producing the most benign fouling and strongest health for the microfragments at that time. The Green Film Summer was second highest and the Clean Start method had the lowest means. Different letters indicate significant differences (α = 0.05), as determined by Dunn’s Multiple Comparison *posthoc* test. Error bars are SEM.

## Discussion

The Green Film method required managing the formation of an algal film throughout the tank to occupy vacant substrate that might otherwise be colonized by noxious foulers prior to placing microfragments into the tank. Pre-selecting green film also resulted in healthier CCA, an important coral settlement inducer, and less nuisance algae present on tank surfaces when microfragments were sampled for this study. Specifically, when compared to the Clean Start method the Green Film method in winter resulted in ∼50% higher CCA health and recruitment scores and a concomitant ∼13% increase in nuisance algae scores (higher scores mean less nuisance algae; Table 1), suggesting that the Green Film might be able to influence the health and rate of tissue expansion in young corals.

**Table 1.** Summary of tank and plate fouling metrics by culture method. Each method was implemented sequentially over 2 years in four culture tanks (230-460L)

| ***Culture Method*** | Months Tested | # Genets | # Frags | # Tanks | Where? | CCA Health* | CCA Recruit* | Nuisance Algae** | Total Score |
| --- | --- | --- | --- | --- | --- | --- | --- | --- | --- |
| *Clean Start* | 4 | 17 | 162 | 6 | Tank | 1.9 | 2.3 | 4.5 | 8.7±0.2 |
|  |  |  |  |  | Plate | 2.1 | 2.3 | 4.2 | 8.0±0.2 |
| *Green Film Summer* | 6 | 29 | 242 | 10 | Tank | 3.1 | 2.4 | 4.1 | 9.7±0.2 |
|  |  |  |  |  | Plate | 3.1 | 2.6 | 4.3 | 10.0±0.2 |
| *Green Film Winter* | 3 | 21 | 121 | 6 | Tank | 3.2 | 3.7 | 5.0 | 11.9±0.9 |
|  |  |  |  |  | Plate | 2.9 | 3.5 | 4.8 | 11.02±0.2 |

\* higher values indicate greater CCA health and recruitment, max 4 points
\*\* higher values indicate lower amount of nuisance algae in tank, max 5 points

The higher performance with growth and fouling metrics of the Green Film Winter may have been facilitated by a number of factors. Although the tanks generally had consistent biophysical parameters throughout the year, the reduced temperature of the incoming seawater prior to system heating and reduced irradiance during the winter months may have slowed the colonization of nuisance algae allowing for more effective eradication using grazers.

In this study all fouling control methods were effective at preventing filamentous algae overgrowing fragments, but cyanobacteria was not equally controlled. Cyanobacteria was present in substantial quantities in the Mini Reef and constantly in need of control in the Clean Start method using grazers and periodic remediation. In contrast, cyanobacteria was nearly absent from the Green Film tanks. Noxious phototrophs, such as cyanobacteria, are suspected to engage in allelopathy and elicit poor microbial ecology (Kuffner and Paul 2004, Vermeij et al. 2009). Kuffner and Paul (2004) found that the presence of the cyanobacteria *Lyngbya majuscula* in coral settlement chambers significantly decreased survival of *Acropora surculosa* larvae and recruitment on unoccupied substrate in *Pocillopora damicornis* (Kuffner & Paul 2004). It is possible that similar deleterious effects may have been occurring in the Mini Reef and Clean Start tanks, as well.

The advantages and disadvantages of these three culture methods for restoration are summarized in Table 2. The Green Film method reduced the mean growth time and was more replicable because it: 1) had an easy startup procedure; 2) created a standardized fouling community; and 3) used active grazer management. Elsewhere, if similar benign algae can be selected for or inoculated in new grow-out tanks it may allow for increased growth rates in microfragment production systems in other parts of the world, enabling more meaningful comparisons between institutions.

**Table 2.** Advantages and disadvantages of each culture method for restoration

| **Culture Method** | **Advantages** | **Disadvantages** |
| --- | --- | --- |
| ***Mini Reef*** | • Familiar strategy<br>• Approximates fouling of a reef<br>• Low maintenance time once established (0.25-0.50 h/tank/day)<br>• Minimizes algal overgrowth of microfragments<br>• No tank restarts needed | • Waiting required– lengthy and variable establishment times (1+ yr)<br>• Slowest grow out time achieved here<br>• Cyanobacteria potentially problematic<br>• Harbors an irreproducible mixture of foulers, potentially resulting in a tank effect |
| ***Clean Start*** | • Easy to start, no waiting<br>• Reduced subset of foulers to manage due to early settlement dynamics<br>• Fouling actively managed<br>• Establishes CCA quickly<br>• Fast grow-out time | • High maintenance time (1-4 h/tank/day)<br>• Threat of nuisance algae bloom is high throughout the tank's life<br>• Restart required in 4-6 months to avoid mini reef conditions |
| ***Green Film*** | • Preselects for benign green algae<br>• Low maintenance time (0.25-1 hour/tank/day)<br>• Simple, replicable fouler management<br>• Fastest grow-out time<br>• Nuisance algae blooms are infrequent | • Two-week waiting period for film to develop<br>• Restart required in 4-6 months to avoid mini reef conditions |

We suggest that the beneficial Green Film method might be a strategy to minimize culture variability and may be key to the widespread implementation and improvement of land based culture for coral restoration and experimentation.

## Supporting information

Supplemental Table 1 and Table 2, and will be used for the link to the file on the preprint site.

## Acknowledgements

The authors would like to thank Tom Moore and Jennifer Moore (NOAA) and Beth Firchau (Association of Zoos & Aquariums) for their comments on an initial draft. This project was funded through, Smithsonian Institution, The Smithsonian’s Women’s Committee, the Paul M. Angell Family Foundation, OceanKind, Revive & Restore, the Scintilla Foundation, the Zegar Family Foundation, the William H. Donner Family Foundation, and the Cedar Hill Foundation. This manuscript has an HIMB citation number of # [add in publication].

