## Supplemental Table 1 and Table 2, and will be used for the link to the file on the preprint site. for "Tank Fouling Community Enhances Coral Microfragment Growth"

**Supplemental Methods and Metrics**

**Methods**

*General Care of the Coral, Microfragmentation and Sequential Husbandry Methods*

Once collected, colonies were housed outdoors in fiberglass tanks fed by filtered seawater at a rate of ~1.5 L/min and supplied with 2-cm air stones for water circulation. Seawater was filtered to 25 µm prior to passing through a 24”-sand filter (Hayward Industries Inc, Newport News, VA) to remove excess detritus, algae, and pest invertebrates, such as *Phestilla sp.*

Tanks were covered with shade cloth and a clear polycarbonate rain sheet in full sun. Sunlight was attenuated to reduce its intensity. PAR values were 200-1000 µmols/m^2^/sec during daylight hours measured by an electronic PAR meter (Apogee Instruments, North Logan, UT). Additionally, each lid was propped open for full sun for ~4 h each day to further regulate incoming light.

Water conditions in the tanks were relatively constant throughout the study in all experimental tanks due to flow-through seawater and the addition of two 500 W heaters used in each tank winter (finnex.net, Inkbird Technical Corporation Limited, Shenzen, China), and a 1.5-hp chiller (model details, ecoplususa.com) used in summer and fall. The temperature in the tanks ranged from 25–27.5 °C throughout the study with winter temperatures having a slightly narrower range (25–26.5 °C) and all other seasons slightly wider range (25–27.5 °C). The salinity and pH were constant throughout the study period at approximately 35 PPT and 8.0, respectively. All tanks received this general care, except during tank set-up and management of the fouling communities which differed between the culture methods.

*Microfragmentation*

Collected colonies were progressively reduced to yield thin ~0.5-cm microfragments according to Page et al (2018). Specifically, fragments were reduced into single branch sections with a hammer and chisel, and then each section was apically bisected with a diamond-bit rotary tool (Robert Bausch Tool Corporation, Mt Prospect, IL) to yield flat slabs of tissue. Next, using a diamond band saw (www.gryphoncorp.com) this material was cut into flat 0.5-cm strips and subdivided into 0.5 cm^2^ pieces. Finally, excess skeleton on the base of each fragment was shaved off to produce a flat section of live coral attached to ~1 mm of skeleton. Once cut, microfragments were glued to a sheet of acetate (Samsill Corporation, Ft Worth, TX) using extra thick super glue gel (Bulkreefsupply.com). Microfragments were evenly spaced such that 30 fragments each from 2 genotypes were mounted to each sheet. The acetate sheet was then clipped to a 0.25-cm-thick, clear acrylic sheet of equal size (Plaskolite, Columbus, OH) with 4 plastic seaweed clips (www.amazon.com). Microfragments were then placed in a flowthrough tank for grow out to yield thinly sheeted tissue.

*Sequential Husbandry Methods*

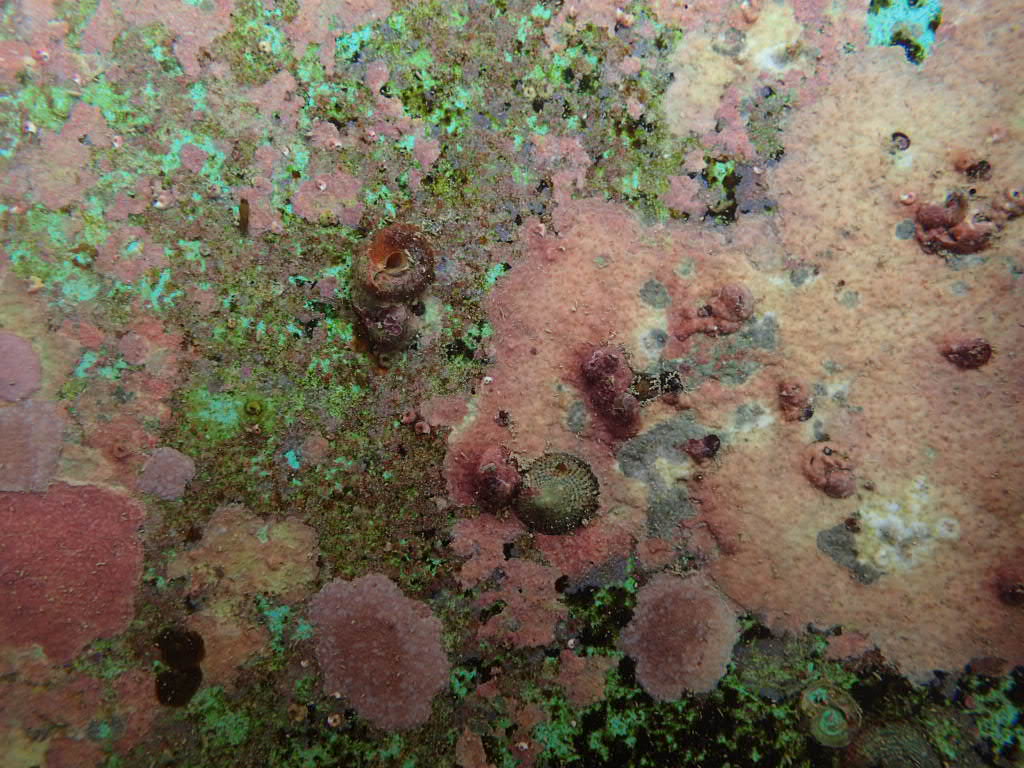

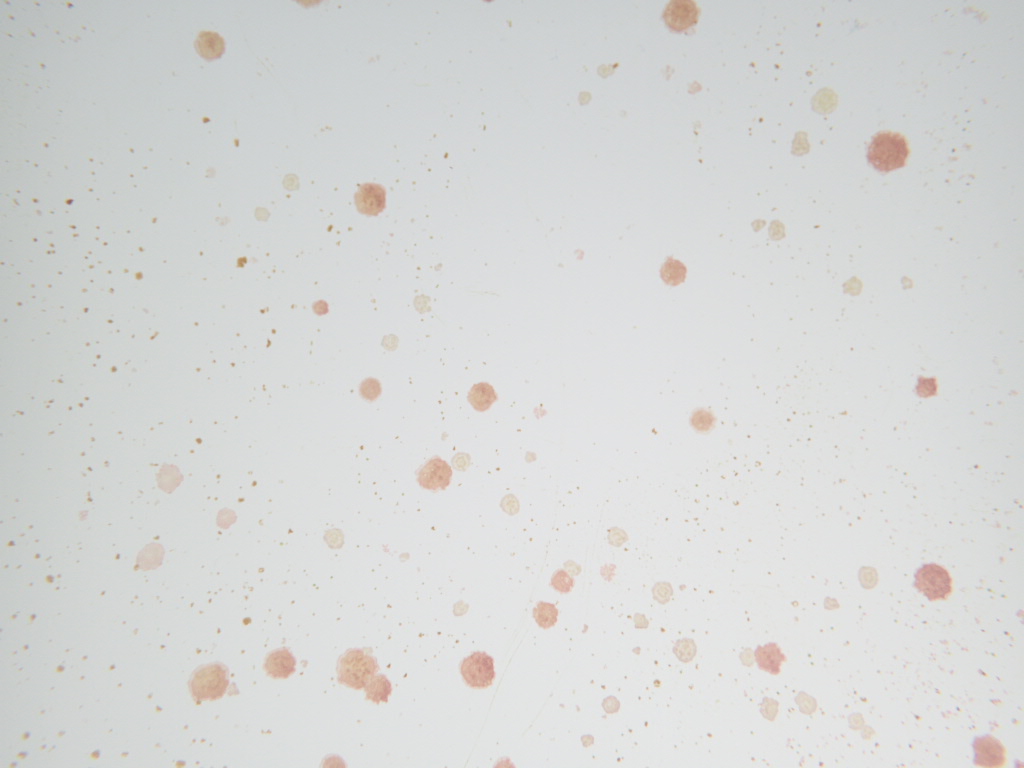

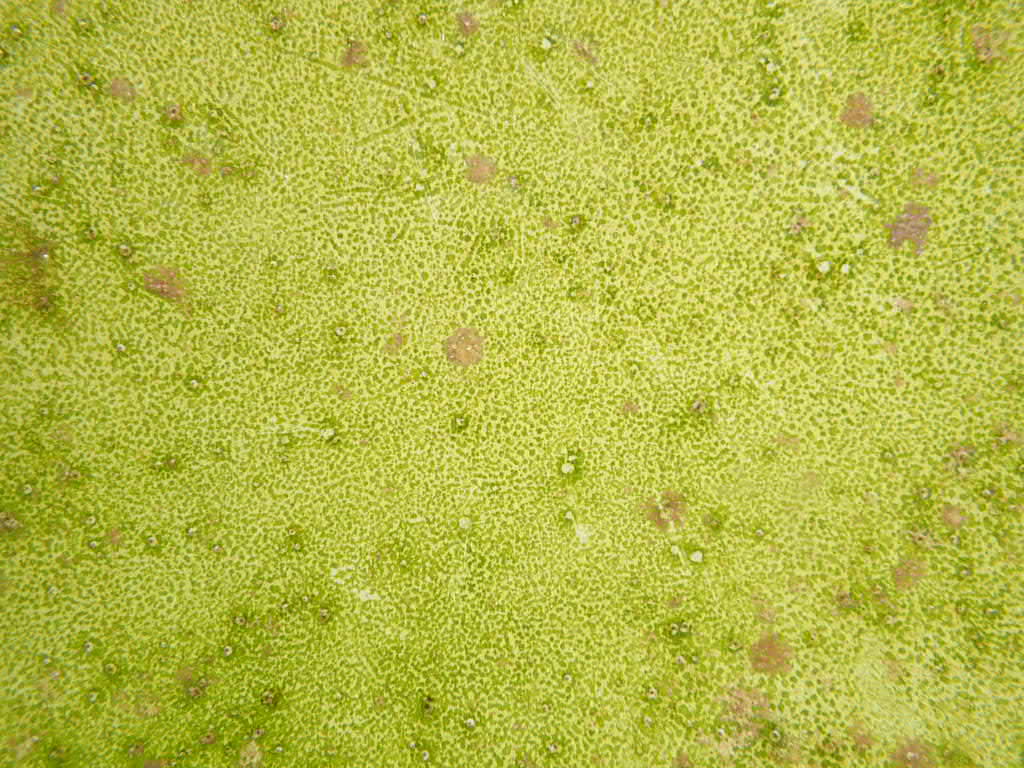

a

b

c

**Fig 1S** Examples of the types of fouling communities typically found on the walls of the tanks of the sequential culture methods, including; a) Mini Reef, b) Clean Start, and c) Green Film

Mini Reef: FEB to MAY 2020. A long established 630-L reef mesocosm equipped with flow through seawater and turbulent flow was retrofitted to grow microfragments. Prior to the addition of the microfragments, the tank was scrubbed, drained, freshwater scoured, and refilled to promote existing CCAs. Live rock, and the grazing fish *Zebrasoma flavescens* were removed and 60 *Trochus intextus* were added to control nuisance algae. Finally, the tank was equipped with aeration and rain guards as described above and siphoned daily to remove excess snail detritus. The resulting fouling community was characterized mainly by thickly encrusted CCAs and cyanobacteria in addition to peysonellid algae, diatoms, and green film algae, with little unoccupied substrate (Fig 1S a). Smothering filamentous algae were notably absent from the vessel. Maintenance time for this method was 0.25–0.50 h/day

Clean Start: MAR 2020 to SEP 2020. In this method, four tanks (230–460 L) were used to grow coral microfragments. They were cleaned to minimize nuisance algae and promote CCA growth. To facilitate this process, tanks were seeded with live CCA chips and 60-80 *Trochus intextus* were added upon startup. Fouler selection occurred during grow-out through a 5-fold process: 1) attaching 4 magnetically-held grazer cages along tank sidewalls, to target diatoms and cyanobacteria (moved daily); 2) replacing fouled seawater spigots, microfragment mounts, and clips (biweekly); 3) laying microfragment plates on the tank floor or caging them overnight with five *T. intextus* to expedite removal of newly settled diatoms and cyanobacteria (as needed); 4) smothering cyanobacteria with epoxy or super glue gel and siphoning up newly settled filamentous algae (as needed); and, 5) temporarily removing microfragments when diatoms and cyanobacteria bloomed in the tank to scrub, drain, freshwater scour, and refill the raceway (1-3 times/month). This method produced a fouling community characterized mainly by thin CCA crusts and uncolonized substrate in addition to small localized patches of cyanobacteria, diatoms and, green film algae (Fig 1S b). Filamentous algae colonization was rare using this method. Maintenance time for this method was 1–4 h per tank per day.

Green Film: SEP 2020 to AUG 2021– In this method four tanks (230-460L) were managed to select for green film algae on the walls. The final culture method required ~2 weeks to curate a benign green film algae in the tanks prior to grow-out of microfragments (Fig 1S c). This fouling community was produced using the following process: 1) prior to the addition of new microfragments, clean, acid-washed tanks were seeded with CCA chips and allowed to accumulate diatomaceous algae for ~1 week; 2) the tank was scrubbed, drained, freshwater scoured, and refilled to remove all algae other than CCA; 3) after an additional week, this process was repeated 1-2 times until scour-resistant green film algae was lightly coating surfaces in the raceway; subsequently, 4) 40-50 grazing *T. intextus* were added to the tanks along with three magnetic grazer cages (each with five snails) and newly cut microfragments (see Fig 2S for details). To facilitate colonization of green film algae tanks were siphoned daily and cages repositioned. Every two weeks seawater spigots, microfragment plates and clips were replaced prevent diatom overgrowth. The resulting fouling community was characterized mainly by green film algae in addition to sparse CCA colonization. Unoccupied substrate was rare using this method. Both filamentous algae and cyanobacteria were nearly absent from grow-out tanks. Maintenance time for this method was 0.25–1 h/tank/day.

**Remediation Week 1**

- Remove CCA rubble to a separate dish with FSW
- Scrub the tank and PVC substrates completely clean of diatoms, then drain, and scour with a direct stream of freshwater for 1 min to promote scrub-resistant green film algae and CCA.
- Replace air stones with clean ones
- Immediately refill the tank with filtered seawater, keeping the exposed walls damp at all times to prevent algae die off.
- Wait an additional week for further algal development

**Remediation Week 2**

- Prior to remediation the tank should contain a mixture of diatoms and a small amount of green film algae
- Remediate the tank once more by draining, scouring, and refilling it
- Immediately add to the tank 40 *T. intextus* that do not have macroalgae or cyanobacteria fouling their shells, and attach 3 clean grazer baskets to the tank walls, each with 5 snails to graze the enclosed area
- Grazer cages: small plastic drawer- organizer baskets with magnets zip-tied at ends, and held in place ends by magnets on tank exterior

**For One Week**

- Prop tank lids open daily (4 h/day) to expose the tank to unattenuated sunlight
- Allow the fouling community to develop unimpeded without the addition of snails or siphoning for 1 week or until noticeable diatomaceous algae is present
- In Hawaii diatomaceous algae appear as delicate tufts of brown algae or as a brown film

**Readying The Tank**

- Scrub and rinse the tank clean of any previous foulers using 3% muriatic acid and freshwater
- Clean PVC fittings for seawater outfalls, overflow strainers, air stones, and snail cages, free of previous foulers in 1:10 dilution of 3% Muriatic acid solution prior to use

**Fill the Tank With FSW**

- Replace air stones, filtered seawater outfalls, and overflow strainers
- Clean and add unfragmented rubble with 100% healthy CCA coverage
- Wash the CCA by:
- Scrubbing the pieces free of detritus using a coarse bristle brush
- Freshwater scouring each piece using a direct stream for 1 minute
- Cover the tank with a plastic lid

**Development of Green Film**

- Siphon and move the cages daily to cover any newly settled diatoms or cyanobacteria, filamentous algae should not be present initially if procedure is performed correctly
- If successful, in ~1–2 weeks light green film algae should become the dominant constituent on tank surfaces
- Add new microfragments as needed

new microfragments can be added as needed

new microfragments can be added as needed

new microfragments can be added as needed

new microfragments can be added as needed

new microfragments can be added as needed

**Fig 2S** Flow chart for creating green film in tanks

**Metrics**

**Table 1S** Specific scoring criteria for tank and microfragment plate metrics. The higher the score for a metric, the healthier the fouling community was considered for the tank or plate.

| CCA Health: the amount of CCA growing or dying in the tank | CCA recruitment: the ratio of new CCA growing in the tank compared to graze resistant competitors | Nuisance Algae: how much algae was throughout the tank? |
| --- | --- | --- |
| 1 - CCA is dying in large sections (area > business card) | 1 - No recruitment of CCA | 1 - Film/Diatoms + heavy hair algae and/or cyanobacteria |
| 2 - CCA is dying in small sections (area < business card) | 2 - Increasing peysonellids/ CCA | 2 - Film/Diatoms + medium hair algae and/or cyanobacteria |
| 3 - CCA is not dying | 3 - No change peysonellids/ CCA | 3 - Film/Diatoms + light hair algae and/or cyanobacteria |
| 4 - Healthy & good CCA growth | 4 - Lower peysonellids/ CCA | 4 - Substantial diatoms and green film algae |
|  |  | 5 - Light diatoms, green film algae, or nothing |

**Table 2S** Specific scoring criteria for the health of the microfragment metrics. The higher the score for a metric, the healthier the microfragment was considered.

| Polyp Extension | Change in Paling/ Tissue Recession | New Tissue Integrity |
| --- | --- | --- |
| 1 - ~25% of frags extended | 1 - 3+ frags are pale or receded | 1 - No growth |
| 2 - ~50% of frags extended | 2 - 1+ frags are pale or receded | 2 - Thin and pale growth |
| 3 - ~75% of frags extended | 3 - 1+ frags are 50%+ paled or receded | 3 - Extra thick and slow growth |
| 4 - ~100% of frags extended | 4 - No new paling or recession | 4 - Thick and fast growth |
|  |  | 5 - Thin and fast growth |
